# Give It Time: Green Anacondas Show Prolonged Behavioral Responses to Enclosure Enhancements

**DOI:** 10.1101/2025.04.14.648616

**Authors:** Maria Inês Poinho, Jacira Carvalho, Gabriela Morello, Francisco Ramos, Ana Magalhães

**Affiliations:** Behavioural Sciences, Instituto de Ciências Biomédicas Abel Salazar, Universidade do Porto (ICBAS), Porto, Portugal; Laboratory Animal Science, (i3S) – Instituto de Investigação e Inovação em Saúde and Instituto de Biologia Molecular e Celular (IBMC), Universidade do Porto, Porto, Portugal; Zoo Santo Inácio, Gaia, Portugal; Addiction Biology, (i3S) – Instituto de Investigação e Inovação em Saúde and Instituto de Biologia Molecular e Celular (IBMC), Universidade do Porto, Porto, Portugal

**Keywords:** *Eunectes murinus*, Environmental enrichment, Captivity, Behavior

## Abstract

Information about green anacondas (*Eunectes murinus*) in human care is limited, especially concerning their behavior, welfare, and overall well-being. To help bridge this knowledge gap, this study evaluated how exhibit design enhancements influence *E. murinus* behavior. The aim was to assess immediate behavioral changes following enclosure modifications and investigate potential habituation over time, ultimately evaluating the effectiveness and longevity of the behavioral changes induced by the improved environment. Systematic observations were divided into three phases: before enclosure improvements, immediately after these modifications, and three weeks later. Enclosure modifications focused on areas less frequented by the anacondas during Phase 1. Results revealed decreased inactivity in Phase 2 and more balanced space use in Phase 3, with anacondas occupying previously underutilized areas, including the terrarium’s vertical dimension. Interaction with exhibit enhancement items increased in the study’s final phase, where a decrease in engagement would be anticipated due to habituation. These results suggested the anacondas required additional time to acclimate to the modifications, even though environmental factors may have contributed to these changes. This study highlights how enclosure enhancements may benefit green anacondas’ welfare while underlining the importance of providing sufficient acclimation time to accurately evaluate the animals’ complete behavioral response.

**Simple Summary:** Green anacondas (*Eunectes murinus*) remain largely understudied, particularly in terms of their behavior and welfare in human care. To address this knowledge gap, improvements were made to the enclosure of green anacondas in captivity, and their behaviors were observed before and after the modifications. The observations were divided into three phases: before the changes (Phase 1), immediately after (Phase 2), and approximately three weeks later (Phase 3). The modifications focused on areas that were rarely utilized during Phase 1. After the improvements, the anacondas became more active and used their enclosure more evenly, including at varying heights. The anacondas interacted more with the new enclosure features after the three weeks, when increased disinterest would have been expected, although environmental conditions may have influenced these changes. These findings show how enclosure modifications may help enhance the welfare of green anacondas and emphasize the need for research that allows enough time for the animals to fully adjust, ensuring a complete understanding of their response.

## 1. Introduction

Green anacondas (*Eunectes murinus*) are predominantly found across South America, where they inhabit various environments, such as forests, savannas, and wetlands (Calderón et al., 2021). As ambush predators that target aquatic prey or animals approaching the water to drink, anacondas have a preference for still water covered in plants (Rivas, 2007; Rivas & Jaremko-Wright, 2023). Neonate green anacondas have often been found in quiet, shallow pools formed in borrow pits, particularly beneath vegetation, twisted around plant bulbs and roots, taking advantage of their coloration for camouflage (Rivas et al., 2016).

During dry periods, as a species adapted to both land and freshwater environments (Calderón et al., 2021), adult anacondas may cluster in riverbank depressions, where they remain mostly inactive, displaying tolerance for close contact with other anacondas until wetter conditions resume (Rivas, 2000 in Rivas et al., 2016; Rivas et al., 2016). In fact, the dry season (January – April) coincides with the mating period (Rivas et al., 2007a; Rivas et al., 2007b), and while lacking a consistent pattern, reproductive activity does not follow a fixed schedule, sometimes continuing for weeks (Rivas et al., 2007a). Rivas (2007) stated that throughout the breeding season, green anacondas – which become sexually mature around the age of three to four years (Miller et al., 2004) – form mating balls, i.e., clusters of intertwined snakes in which multiple males compete for the opportunity to mate with a single female – suggesting that green anacondas exhibit some level of social interaction, at least during this period of the year.

Although additional studies may be needed for full confirmation, according to Rivas et al. (2007a), *E. murinus* activity levels appear to vary according to the environment’s temperature, decreasing during the hottest parts of the day. This behavioral change was particularly evident to the authors on cloudy days, with anacondas exhibiting overall increased activity, further reinforcing that activity levels were related to temperature rather than time of day (Rivas et al., 2007a). Indeed, since snakes are ectotherms – and therefore their body temperature fluctuates with environmental conditions, relying on external sources for regulation – they depend on their behavior to ensure they maintain suitable body temperatures (Sunday et al., 2014). Nonetheless, while anacondas tended to be more active during the evening and nighttime, Rivas et al. (2007a) observed that they engaged in activities such as moving, foraging, constricting prey, and mating throughout both day and night, with basking behavior commonly observed in pregnant females. Similarly, green anaconda neonates were observed engaging in various activities – sheltering under vegetation, moving, basking, buried or in caves, under water, and eating – though the specific timing of these behaviors was not documented (Rivas et al., 2016).

Although the last assessment for the International Union for Conservation of Nature (IUCN) Red List of Threatened Species, in 2014, classified *E. murinus* as “Least Concern”, the current population trend is unknown, and the species faces several threats (Calderón et al., 2021). The growth of beach tourism and habitat disturbance (Smith et al., 2016), as well as hydroelectric dams causing the destruction and deterioration of certain aquatic and riparian habitats (Calderón et al., 2021), might pose challenges to its conservation. Moreover, anaconda skins have been seized in the past (Rivas, 2007) presumably for leather production, and in certain regions *E. murinus* fat is used for medicinal purposes (Alves et al., 2007). In addition to these factors, green anacondas are feared by the population since they are the heaviest snakes in the world (Calderón et al., 2021) and, according to a 2016 study, the larger the anaconda the bigger the probability of it being killed, possibly as a “preventive or punitive killing” based on fear (Miranda et al., 2016, p. 50).

While reptiles, including green anacondas, are often met by most people with apprehension, leading to the misconception that as “cold-blooded” animals they lack feelings, research shows that reptiles can experience emotions and demonstrate cognitive abilities (Lambert et al., 2019). However, studies regarding reptilian behavior, welfare, and overall well-being are not exactly abundant, at least compared to other taxa (Rosenthal et al., 2017; Binding et al., 2020). If we narrow the focus to green anacondas under human care, a literature search revealed no published studies specifically addressing this topic. To help bridge this information gap, this study investigated the impact of terrarium improvements on the behaviors and space utilization of a pair of green anacondas in captivity. To achieve this, the following objectives were established: A) assess the anacondas’ behavior and use of space before any enclosure modifications, B) examine the immediate changes in behavior and space utilization following terrarium enhancement and C) evaluate potential habituation approximately three weeks after the enclosure modifications, assessing the effectiveness and longevity of the behavioral and space utilization changes prompted by the enhanced environment.

## 2. Materials and Methods

### 2.1. Animals and husbandry

This research occurred at Zoo Santo Inácio (41°05’34.4”N, 8°32’15.5”W), the largest zoological park in northern Portugal, between January and June of 2024. Two captive-bred adult green anacondas (*Eunectes murinus*), one male and one female, were observed in this study. At the time of the research, both animals were between 16 and 18 years old and had been housed in the zoo since 2008. According to the latest measurements and weightings – in 2018 and 2023, respectively – the male anaconda measured 286 centimeters (total length) and weighed 29 kilograms, while the female was slightly smaller at 256 centimeters (total length) and 20 kilograms. The distinction between the two anacondas was based on the female being smaller and having a lighter coloration than the male. Each anaconda was fed weekly with a single dead rabbit or guinea pig and, occasionally, a pair of dead quails, placed in the terrarium with a long-handled tong. There was no fixed schedule for enclosure cleaning, but it was performed at least once a week. Since being cared for by the zoo, the anacondas have never reproduced nor had contact with other animals of their species apart from each other. During this study, the female anaconda underwent ecdysis over approximately 11 days, between the 11^th^ and 22^nd^ of May 2024.

### 2.2. Behavioral observations

#### 2.2.1. Preliminary observations

*Ad libitum* observations were conducted before starting systematic data collection to develop the species’ ethogram, from the 8^th^ to the 25^th^ of January 2024. These observations, totaling 20 hours, were distributed between morning and afternoon sessions. In total, 22 behaviors were identified (see *Figure S1* for a detailed description and photographic examples) and were organized into different behavioral categories (*Table 1*).

**Table 1.** Green anaconda’s behavioral categories, descriptions, and the specific behaviors in each category.

| Behavioral category | Description | Included behaviors |
| --- | --- | --- |
| Inactive | The anaconda is motionless | Resting submerged in water<br>Resting on land |
| Locomotion | Movement between locations using lateral body undulations | Entering the lake<br>Exiting the lake<br>Swimming<br>Slithering |
| Feeding | Related to food detection and ingestion | Food pre-capture<br>Food capture |
| Climbing | The anaconda moves up or down to reach different heights | Climbing up<br>Climbing down |
| Body maintenance | Related to physiological and physical needs | Yawning<br>Rubbing |
| Interaction with transparent boundaries | Engaging with the enclosure's high windows | Swimming against the high window<br>Crawl up the high window |
| Social interaction | Involving interactions with conspecifics | Coiling around conspecific<br>Proximity to a conspecific |
| Exploration | Performed to explore the environment | Tongue flicking<br>Head or body elevation<br>Head or body lowering |
| Engaging with habitat enhancement items | Involving interactions with items used for habitat enhancement | Resting on habitat enhancement items<br>Rubbing against habitat enhancement items<br>Slithering on habitat enhancement items |
| Not visible | The anaconda is out of sight |  |

#### 2.2.2. Systematic observations

The systematic observations were divided into three distinct phases (*Figure 1*). The first phase, conducted before any modifications to the terrarium, lasted nine consecutive days, during which data on the anacondas’ behavior was collected. This was followed by a one-day interval to implement improvements to the terrarium. The second phase, also lasting nine consecutive days, focused on documenting the animals’ interactions with the improved environment. A 15-day interval was introduced between Phase 2 and Phase 3 to allow the anacondas time to acclimate to the environmental modifications. Phase 3, comprising another nine days of behavioral data collection, aimed to observe the anacondas’ behavior following this acclimation period in order to detect possible loss of interest in the improved areas.

**Figure 1.**
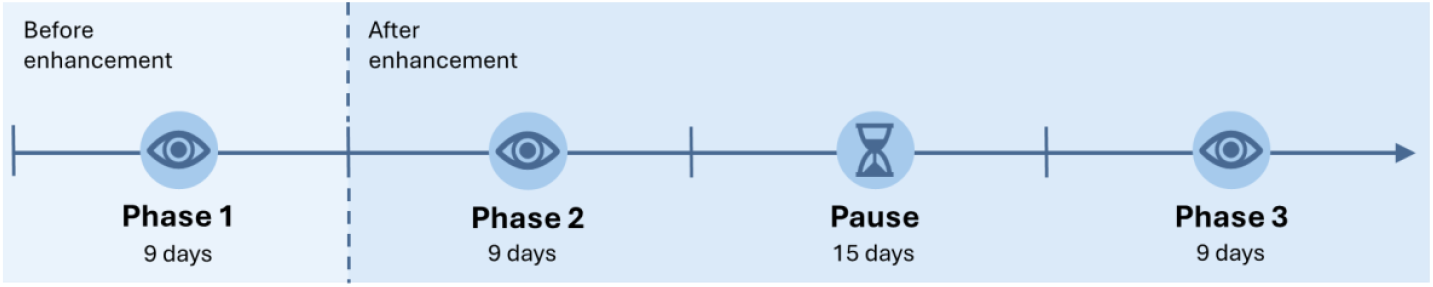
Timeline illustrating the division of the study’s behavioral observations into three distinct phases. Phase 1 occurred before enclosure enhancements, while Phases 2 and 3 occurred after making improvements to the terrarium. A 15-day pause in behavioral data collection occurred between Phase 2 and Phase 3 to allow the anacondas to acclimate to enclosure modifications and assess any potential decline in engagement with the enhanced areas.

Direct observations were carried out between April 29^th^ and June 7^th^ of 2024, with six 30-minute observations each day - three in the morning and three in the afternoon. Morning observations were performed from 10h30 AM to 1 PM and afternoon observations from 2 PM to 4h30 PM, with a 30-minute break between observations in both the morning and afternoon sessions. An instantaneous sampling method was used for all observations (Altmann, 1974). To habituate the anacondas to the observer’s presence, the observer stood in front of the terrarium’s high window for 10 minutes before each morning and afternoon session started.

### 2.3. Study area

Both anacondas were housed in an L-shaped terrarium (approximately 44 m^2^) featuring a high window running along its length, providing a clear view of the entire area (*Figure 2*). A lake approximately 50 centimeters deep was present at the terrarium’s center, and the enclosure’s floor was covered with pebbles and rocks of different sizes (*Figure 2*). The terrarium had irregular and rough walls, appropriate for climbing, as well as elevated artificial rocks with heat lamps directed at them and UV lights placed throughout the space (*Figure 2*). The extremities of the terrarium had different environmental conditions, with an average temperature of 28ºC and a relative humidity of 89% on the left side, and an average temperature of 26ºC and a relative humidity of 93% on the right side. The terrarium was divided into five distinct areas (*Figure 2*) and for all phases the anacondas’ behaviors, locations within the terrarium, and the heights they were at when inactive – categorized as low (≤10 cm), medium (0.11 to 0.80 cm), and high (≥0.81 cm) – were documented every 30 seconds.

**Figure 2.**
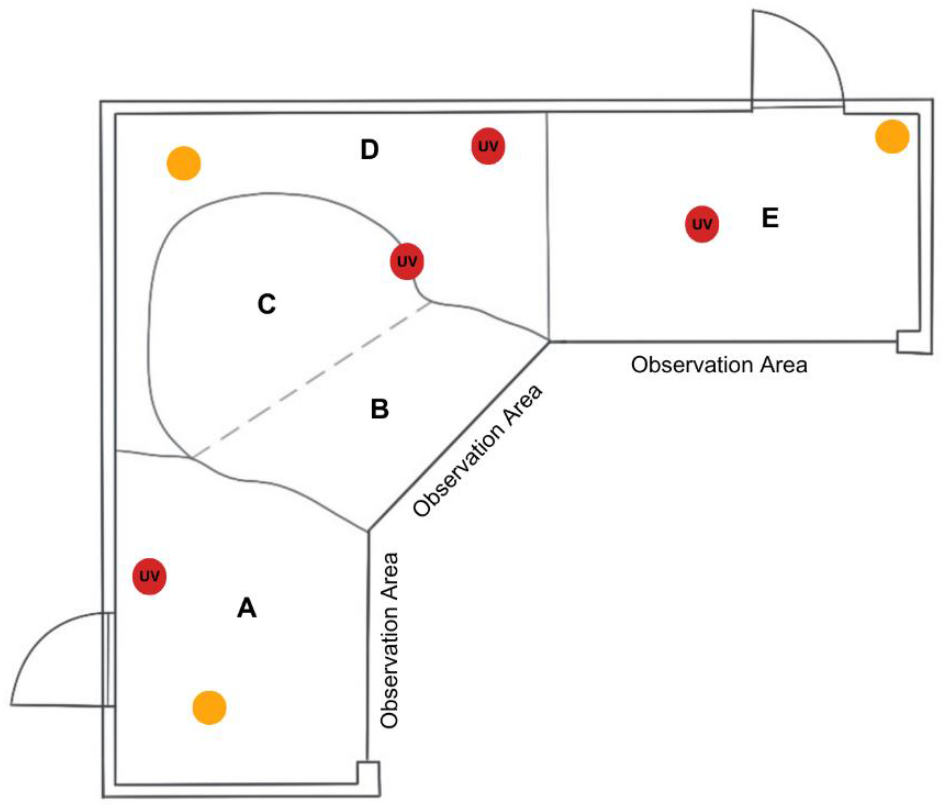
Schematic illustration of the green anacondas (*Eunectes murinus*) terrarium, divided according to the study areas. **(A)** Left side of the terrarium; **(B)** Outer lake; **(C)** Inner lake; **(D)** Center of the terrarium; **(E)** Right side of the terrarium. Quarter circles represent the entrances to the terrarium used by zoo staff. Yellow and red circles mark the location of heat and UV lamps, respectively. Visitors could view the anacondas from dedicated observation areas.

### 2.4. Environmental improvements

Environmental enhancements were designed to offer the anacondas greater choice by incorporating diverse textures and hiding spots. These elements aimed to stimulate natural exploratory behaviors by presenting the snakes with novel sensory experiences and spatial complexity. The environmental improvements were specifically targeted at areas that the anacondas frequented less often during Phase 1 (*Table 2*). As such, the terrarium floor was covered with pine tree bark, a trunk, rocks, and artificial plants. A trunk was placed within the terrarium’s lake, crossing it from one side to the other, with one end submerged and the other end at the lake’s edge, surrounded by artificial plants (*Figure 3*). Additionally, artificial plants were fixed to rocks at the bottom of the lake, creating the effect of plants rising toward the surface (*Figure 3*). The pine bark, trunks, and rocks were repurposed from materials already available at the zoo, while the artificial plants were purchased from a local store.

**Table 2.** Selected items for environmental improvement and their placement within the terrarium.

| Item | Placement | Zone<sup>1</sup> |
| --- | --- | --- |
| Pine tree bark | Covering the terrarium’s floor. | A and E |
| Tree trunks | Across the lake, with one side submerged and the other at the lake’s edge. On the terrarium’s floor. | B and E |
| Rocks | On the terrarium’s floor. | A |
| Artificial plants | Attached to rocks at the bottom of the lake, and around the trunk at the lake’s edge. On the terrarium’s floor. | A, B and E |

<sup>1</sup> Considering *Figure 2*.

**Figure 3.**
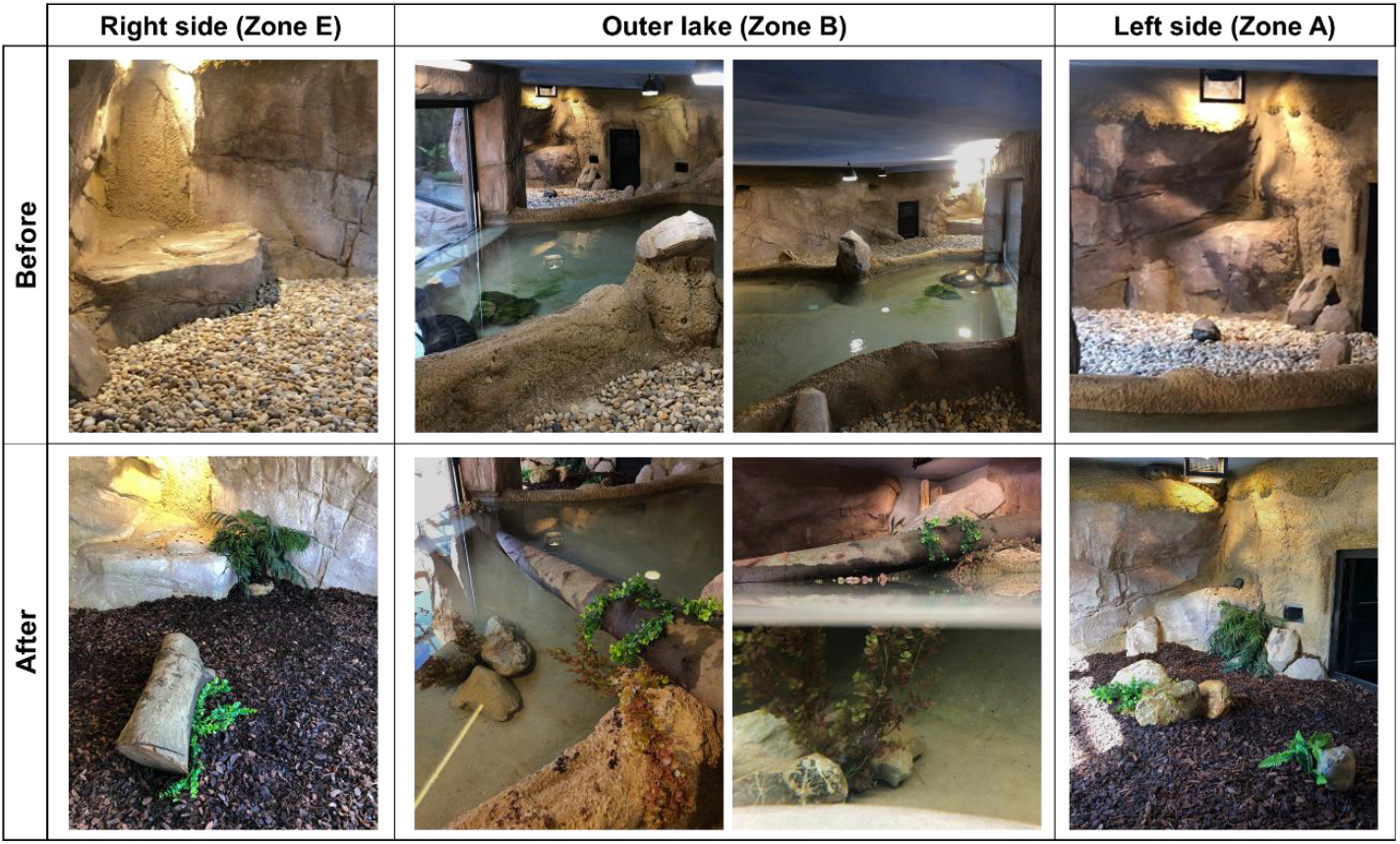
Comparison of the anacondas’ terrarium before and after enclosure modifications. Enhancements included the addition of pine bark, trunks, rocks and artificial plants, and were focused on the right and left sides of the exhibit, as well as the outer lake (i.e., the areas less frequented at the start of the study). The terrarium’s central area and the inner part of the lake remained unchanged.

### 2.5. Statistical analysis

The anacondas’ behavior was evaluated by comparing each behavioral category across the three phases of the study. As the data did not meet the normality and homogeneity assumptions, a Friedman test was conducted with a two-stage linear step-up procedure of Benjamin, Krieger and Yekutieli correction for multiple comparisons. A Wilcoxon test was used to compare engagement with habitat enhancement items between short-term (Phase 2) and long-term (Phase 3) periods since the Shapiro-Wilk test revealed a non-normal data distribution. The significance set for both tests was *p*≤0.05. The statistical analysis of the previous tests was performed with GraphPad Prism version 8.0 (California, USA).

The anacondas’ space occupation was analyzed across the three phases: before enclosure modifications (Phase 1), immediately after enhancements (Phase 2), and approximately three weeks after enclosure modifications (Phase 3). The occupancy frequency was calculated for each of the five zones (*Figure 2*) using the formula: Space occupation (%) = (Zone occupancy / Total occupancies across all zones) x100. Additionally, the number of times a specific snake was seen in a specific area per scan-sampling event (snake sightings) had a binomial distribution and was coded into minimal area use (i.e. ≤10 sightings) or frequent use (> 10 sightings).

The probability of frequent use of each of the areas (response variables) was modeled as a function of time-point of data collection (6-level categorical fixed effect), phase (3-level categorical fixed effect), sex (2-level categorical fixed effect, and the interaction between phase and sex. Procedure GLIMMIX was used on SAS (2018 University Edition, SAS Institute Inc., USA), considering a binomial distribution and a logit link. A two-way ANOVA test performed with GraphPad Prism version 8.0 (California, USA) was used to evaluate the anacondas’ preference for different enclosure levels (low, medium, and high) during inactive periods across all three study phases. Since the data violated normality and homogeneity of variance assumptions, only differences with p≤0.01 were considered statistically significant, with a two-stage linear step-up procedure of Benjamin, Krieger and Yekutieli correction for multiple comparisons.

Relative humidity and heat lamp temperature measured at both the right and left sides of the terrarium (*Figure 2*) were regressed as functions of phase. Heat lamp temperature difference between the right and left sides of the terrarium was also regressed as a function of phase. Procedure GLM was used on SAS (2018 University Edition, SAS Institute Inc., USA), considering a gamma data distribution. Least-squares means were compared considering a 95% confidence interval with a Tukey-Kramer adjustment for multiple pairwise comparisons.

The behavioral category “Feeding” was excluded from statistical analysis as observations never coincided with feeding times.

## 3. Results

### 3.1. Behavioral Analysis

Terrarium modifications appeared to have an impact on *E. murinus* behavior, leading to a lower frequency of inactive behaviors in Phases 2 and 3 (*Figure 4a*), with a statistically significant difference observed in Phase 2 (p=0.036) compared with Phase 1. Social interactions peaked significantly during Phase 2 relative to Phase 1 (p=0.001) but declined to their lowest frequency in Phase 3 (p<0.0001; *Figure 4b*). In contrast, locomotion was significantly higher in Phase 3 compared to both Phase 1 (p=0.021) and Phase 2 (p=0.004; *Figure 4c*). Exploratory behaviors followed a similar pattern, with significantly higher levels in Phase 3 than in Phases 1 and 2 (p=0.041 and p=0.001, respectively; *Figure 4d*). Interactions with habitat enhancement items were also most frequent in Phase 3, surpassing those in Phase 2 (W=618; p<0.0001; *Figure 4e*).

**Figure 4.**
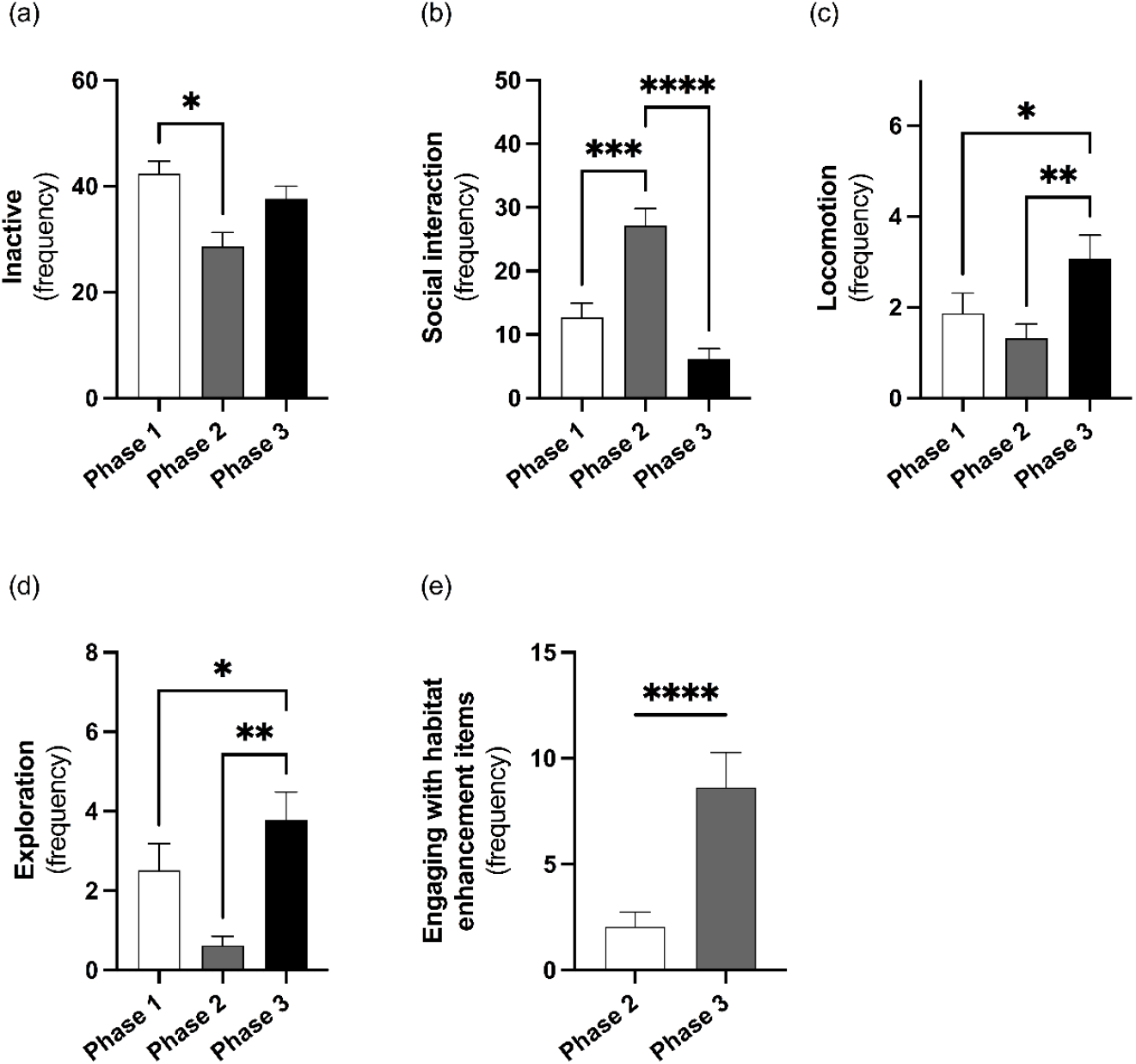
Impact of enclosure enhancement on *E. murinus* behavior across study phases. Panels display the frequency of **(a)** inactivity, **(b)** social interactions, **(c)** locomotion, **(d)** exploration, and **(e)** engagement with habitat enhancement items. **Phase 1 –** Before enclosure modifications; **Phase 2 –** Immediately after enclosure modifications; **Phase 3 –** Approximately three weeks after enclosure modifications. Data is represented as mean ± SEM. *p ≤ 0.05, **p ≤ 0.01, ***p ≤ 0.001, ****p ≤ 0.0001.

No statistically significant differences were observed across phases for “Climbing”, “Body maintenance”, “Interaction with transparent boundaries”, and “Not visible” behavioral categories. Nevertheless, climbing behaviors were the lowest during Phase 2 before reaching their peak during Phase 3 (*Figure S2a*). Body maintenance behaviors were higher in Phases 2 and 3 compared to baseline (Phase 1; *Figure S2b*). Interactions with transparent boundaries showed a progressive increase from Phase 1 through Phase 3 (*Figure S2c*). Instances where the anacondas were not visible occurred only during Phase 1.

### 3.2. Space Occupation and Environmental Conditions

During Phase 1, the male anaconda primarily occupied the inner lake (Zone C: 41.64%) and the terrarium’s center (Zone D: 41.98%), which were used almost equally (*Figure 5*). The male utilized less frequently the terrarium’s left side (Zone A: 9.01%) and outer lake (Zone B: 6.85%), while rarely occupying the terrarium’s right side (Zone E: 0.52%). In this phase, the female exhibited a preference for the inner lake (Zone C: 71.67%), with limited use of the terrarium’s center (Zone D: 16.67%) and outer lake (Zone B:11.67%). The female did not utilize Zones A and E during Phase 1 (*Figure 5)*.

**Figure 5.**
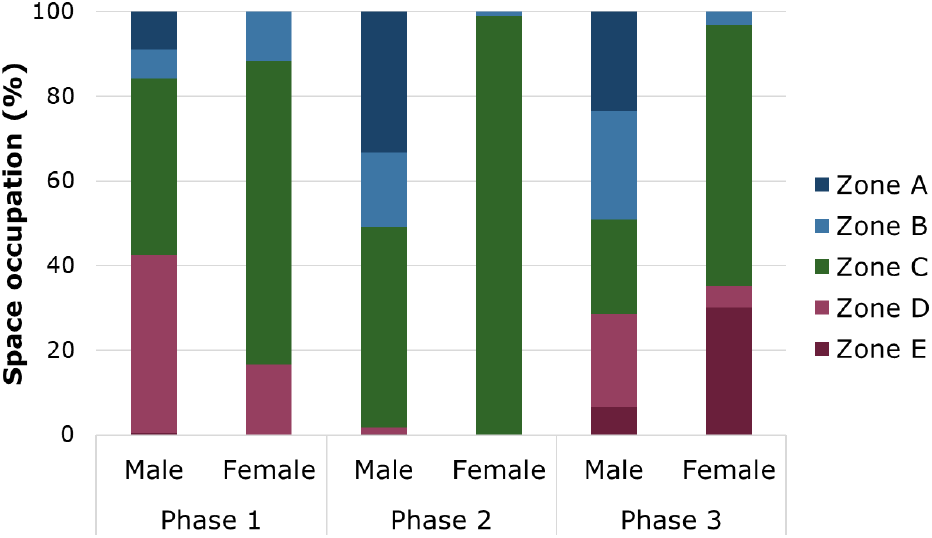
Percentage of space occupation of male and female *E. murinus* across the three study phases. **Zone A –** Left side of the terrarium; **Zone B –** Outer lake; **Zone C –** Inner lake; **Zone D –** Center of the terrarium; **Zone E -** Right side of the terrarium. **Phase 1 –** Before enclosure modifications; **Phase 2 –** Immediately after enclosure modifications; **Phase 3 –** Approximately three weeks after enclosure modifications.

In Phase 2, a phase effect was observed (F_(2, 159)_=3.56; p=0.0306) in the terrarium’s left side (Zone A) usage, which increased significantly compared to Phase 1 (p=0.0231; *Figure 6a*). Since the female anaconda did not utilize Zone A during this phase, data analysis for this zone was limited to the male anaconda, whose occupation of Zone A increased to 33.33% (*Figure 5*). The usage of the terrarium’s outer lake (Zone B) showed significant effects related to sex (F_(1, 318)_=12.02; p=0.0006) and the interaction between phase and sex (F_(2, 318)_=3.92; p=0.0208). Although no significant differences were observed between Phases 1 and 2 or between sexes, the male anaconda occupied this space more than the female, with usage rates of 17.62% and 0.99%, respectively (*Figure 5*). During Phase 2, the inner lake (Zone C) was the most utilized area by both anacondas, with a significant increase in usage compared to Phase 1 (p=0.0098; *Figure 6c*). Statistical analysis revealed significant effects of both phase (F_(2, 320)_=10.96; p<0.0001) and sex (F_(1, 320)_=39.23; p<0.0001) on Zone C occupation. Notably, usage of Zone C differed between sexes, with the male occupying it 47.31% of the time, compared to 99.01% for the female (*Figure 5*). Throughout Phase 2, the terrarium’s center (Zone D) was exclusively utilized by the male anaconda. Usage of Zone D significantly decreased compared to Phase 1 (p=0.0003; Figure 6d), with the male occupying it only 1.36% of the time (*Figure 5*). For Zone D, significant phase (F_(2, 320)_=9,22; p=0.0001) and sex (F_(1, 320)_=18.82; p<0.0001) effects were observed, influencing its occupation. Regarding the terrarium’s right side (Zone E), although a significant phase effect was observed (F_(2, 320)_=10.28; p<0.0001), no significant sex differences in usage were found during Phase 2 (*Figure 6e*). Indeed, the male anaconda showed minimal occupation of Zone E at 0.37%, while the female avoided this zone completely (*Figure 5*).

**Figure 6.**
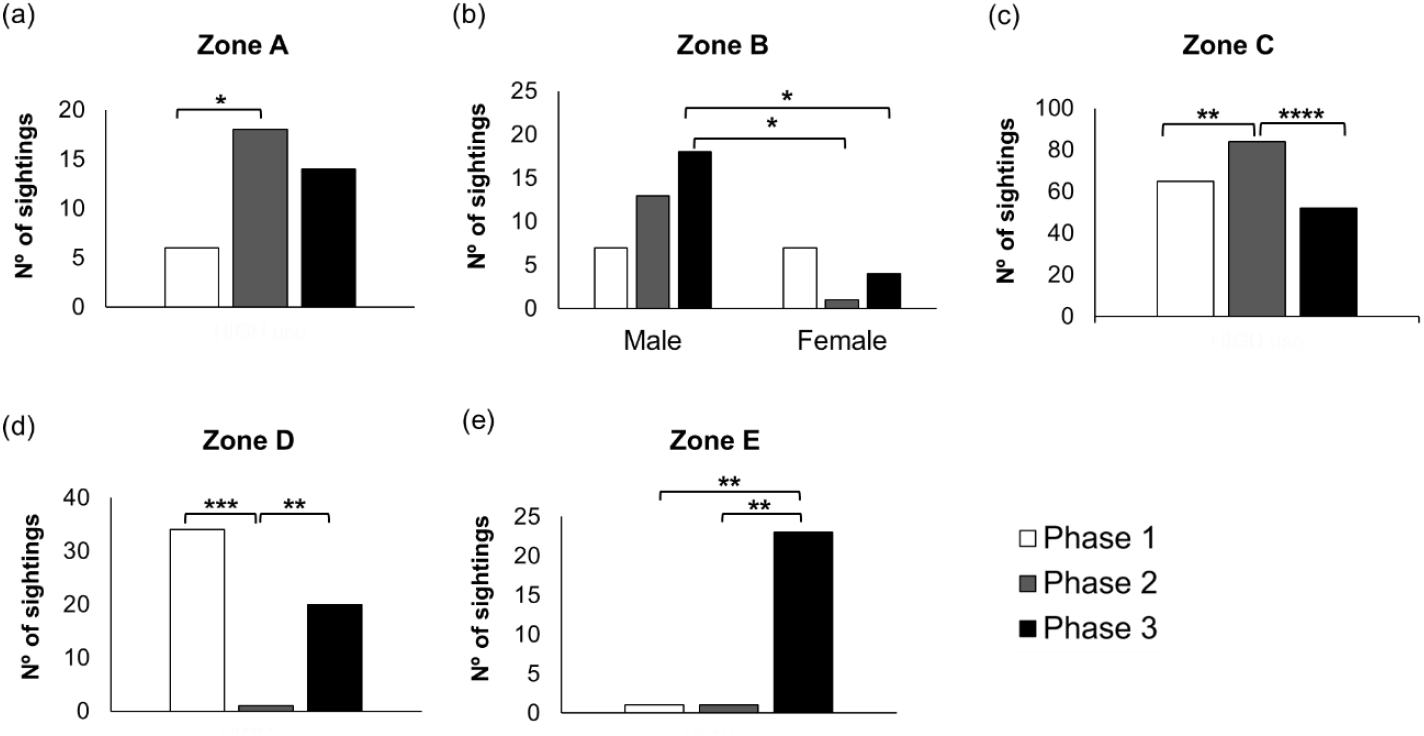
Frequent enclosure use (> 10 sightings) by zone across phases. (**a)** Left side of the terrarium; **(b)** Outer lake; **(c)** Inner lake; **(d)** Center of the terrarium; **(e)** Right side of the terrarium. **Phase 1 –** Before enclosure modifications; **Phase 2 –** Immediately after enclosure modifications; **Phase 3 –** Approximately three weeks after enclosure modifications. Graph **(b)** shows Zone B utilization separately by sex due to a statistically significant interaction between phase and sex. All other graphs present combined male and female zone utilization patterns, depicting the actual number of sightings obtained per snake per zone. *p ≤ 0.05, **p ≤ 0.01, ***p ≤ 0.001, ****p ≤ 0.0001, indicating statistically significant differences between least-squares means from the statistical models.

In Phase 3, analysis of Zone A use was again limited to the male anaconda, as the female continued to avoid it entirely (*Figure 5*). The male anaconda occupied this area 23.58% of its time (*Figure 5*), showing no significant differences compared to previous phases (*Figure 6*). On the contrary, significant differences were detected between the female’s Zone B usage during Phase 2 and the male’s during Phase 3 (p=0.0239; *Figure 6b*), as well as between both sexes’ Zone B usage within Phase 3 (p=0.0268; *Figure 6b*). During this phase, the male occupied Zone B 25.46% of the time, compared to the female’s usage of 3.06% (*Figure 5*). The inner lake (Zone C) was significantly less utilized by both anacondas during this phase (p<0.0001; *Figure 6c*) compared to Phase 2, with the male decreasing its use to 22.28% and the female to 61.64% (*Figure 5*). A notable shift occurred in the terrarium’s center (Zone D) with a significant increase in use by both anacondas compared to Phase 2 (p=0.0050; *Figure 6d*). The male increased its use to 22.19% and the female to 5.19% (*Figure 5*). Phase 3 was also characterized by a significant increase in the use of the terrarium’s right side (Zone E) compared to Phases 1 and 2 (p=0.0031 for both; *Figure 6e*). Space occupation rates increased to 6.48% for the male and to 30.12% for the female (*Figure 5*).

Regarding vertical enclosure use, results showed that there was a significant effect of enclosure height (F_(2, 321)_= 25.42; p<0.0001), a phase effect (F_(1.732, 546.4)_= 5.396; p=0.0071), and an interaction between phase and height (F_(4, 631)_ = 24.45; p<0.0001). During Phases 1 and 2, the anacondas predominantly used high surfaces, with this preference being significantly more pronounced in Phase 1 compared to Phase 3 (p=0.001, *Figure 7*). Low and medium surfaces were rarely utilized during Phases 1 and 2 (*Figure 7)*. In contrast, Phase 3 revealed a more distributed vertical enclosure use, with the anacondas occupying all three levels, although they exhibited a significant preference for the middle level compared to Phases 1 and 2 (p<0.0001; *Figure 7*).

**Figure 7.**
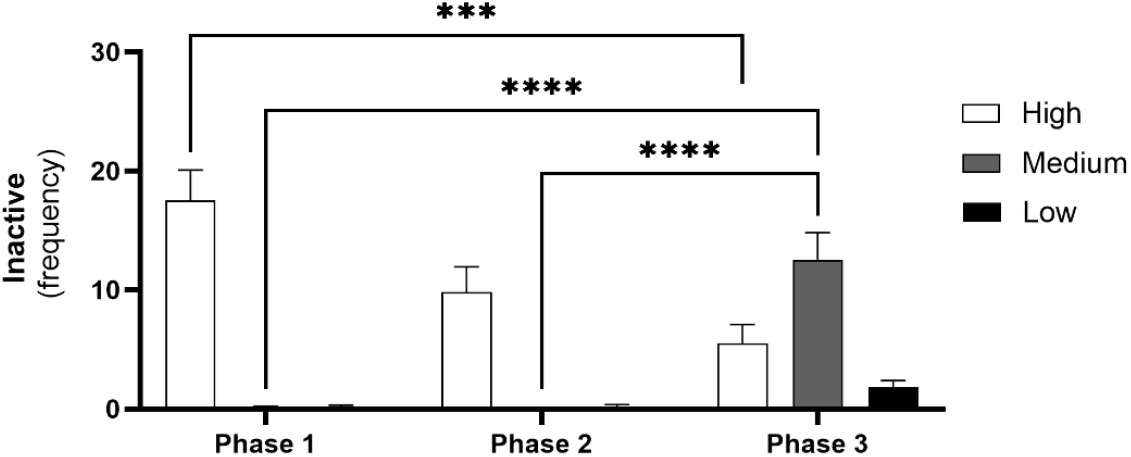
Anacondas’ vertical habitat preference during inactive periods across the study phases. **Low** ≤10 cm; **Medium -** 0.11 to 0.80 cm; **High** ≥0.81 cm. **Phase 1 –** Before enclosure modifications; **Phase 2 –** Immediately after enclosure modifications; **Phase 3 –** Approximately three weeks after enclosure modifications. Data is represented as mean ± SEM. ***p ≤ 0.001, ****p ≤ 0.0001.

Statistical analysis revealed significant differences in temperature across phases and between the left and right sides of the terrarium (*Figure 8a*). Temperatures on both sides followed a similar pattern: decreasing from Phase 1 to Phase 2, then increasing slightly from Phase 2 to Phase 3 (*Figure 8a*). On the left side, statistically significant differences were observed between all phases: Phase 1 and Phase 2 (p<0.0001; *Figure 8a*), Phase 1 and Phase 3 (p<0.0001; *Figure 8a*), and Phase 2 and Phase 3 (p=0.0105; *Figure 8a*). Similarly, the terrarium’s right side exhibited significant differences between all phases: Phase 1 and Phase 2 (p<0.0001; *Figure 8a*), Phase 1 and Phase 3 (p<0.0001; *Figure 8a*), and between Phase 2 and Phase 3 (p=0.0003; *Figure 8a*). Despite these temperature variations, statistically significant differences in the temperature gradient were only found when comparing Phase 1 to Phase 2 (p<0.0001; *Figure 8b*) and Phase 1 to Phase 3 (p=0.0013; *Figure 8b*), with no significant differences detected between Phases 2 and 3. Additionally, data analysis showed that temperature and relative humidity were significantly inversely correlated (p<0.0001).

**Figure 8.**
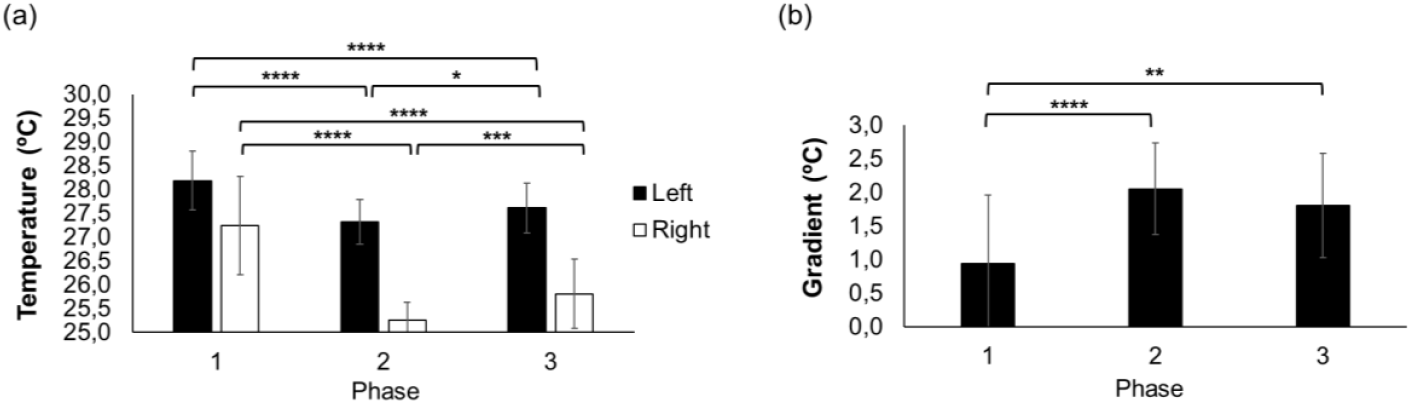
Environmental conditions across phases on the terrarium’s left and right sides. **(a)** Temperature measurements (ºC); **(b)** Temperature gradient (ºC). **Phase 1 –** Before enclosure modifications; **Phase 2 –** Immediately after enclosure modifications; **Phase 3 –** Approximately three weeks after enclosure modifications. *p ≤ 0.05, **p ≤ 0.01, ***p ≤ 0.001, ****p ≤ 0.0001, indicating statistically significant differences between least-squares means from the statistical models.

## 4. Discussion

This research aimed to evaluate changes in *E. murinus* behavior in response to enclosure enhancements and determine whether they preferred the modified areas over the unaltered ones by analyzing their space occupation throughout the process. In doing so, we also present, to the best of our knowledge, the first ethogram of this species in captivity, providing a structured framework for understanding its behavioral repertoire in managed environments.

### 4.1. E.murinus Behavioral Responses to Terrarium Enhancement

Results showed that terrarium enhancements led to the reduction of inactive behaviors, although this difference was only significant for Phase 2. There is no available information on the amount of time *Eunectes murinus* spends on activity or inactivity, either in the wild or in captivity. Although inactivity is often associated with negative affective states – such as remaining motionless in response to threats, poor health conditions, or depression and boredom – it can equally indicate the opposite (Fureix & Meagher, 2015). Given the anacondas’ semiaquatic and ambush-predator lifestyle (Rivas et al., 2007a), and considering the evidence that some reptile species spend around 67% to 98% of their time completely stationary (McElroy et al., 2011), it is likely that prolonged periods of inactivity play a crucial role in their energy conservation strategy. Moreover, as a semi-aquatic species (Calderón et al., 2021) *E. murinus* is frequently observed in its native habitat submerged in water, hidden by surrounding vegetation, while waiting for prey that approaches to drink (Rivas, 2007; Rivas et al., 2007a; Rivas et al., 2016). Therefore, this species’ tendency to remain inactive in an aquatic environment is not unusual, as this behavior aligns with its natural hunting strategy. This is consistent with the idea that periods of inactivity may signal positive affective states, such as the stillness following copulation or feeding, and during sun-basking (Fureix & Meagher, 2015).

Basking contributes to physiological processes such as vitamin synthesis (Ferguson et al., 2003; Karsten et al., 2009), but its most well-known function is thermoregulation (e.g., Smith, 1979; Huey et al., 2012; Stanton-Jones et al., 2018). In ectotherms, behavioral strategies have a bigger impact on body temperature regulation than physiological processes (Stevenson, 1985). For most ectothermic animals, adjusting behavior to different environmental conditions is critical for survival, as solely relying on physiological thermal tolerance is often insufficient (Sunday et al., 2014). Although basking is a common behavioral strategy among reptiles (Brattstrom, 1965), the anacondas in this study were never observed doing so. However, despite a gradual decline over time, the anacondas frequently utilized higher levels of the enclosure (*Figure 7*), which were positioned closest to the heat lamps. On the one hand, the warmth from the lamps may have provided a sensation of thermal comfort similar to sunlight exposure, potentially explaining the anacondas’ preference for remaining inactive on elevated surfaces, suggesting that their inactivity in these areas may have been a comfort behavior. On the other hand, moving closer to the heat lamps may have been a sign of behavioral fever, where “healthy but ‘stressed’ reptiles” search for warmer environments until their stress levels decrease (Warwick et al., 2013, pp. 130). Understanding these inactivity/activity patterns in anacondas could provide valuable insights into their ecology and welfare in managed environments.

During Phase 2, social interactions occurred with notable frequency and may have been related to the female’s skin-shedding process and the consequential release of chemical cues. Reptiles rely on chemical communication for numerous functions, including locating food sources, detecting potential predators, identifying species and individuals, selecting mates, signaling danger, and establishing territory (Mason & Parker, 2010). In snakes, pheromones have been linked to sexual attractiveness, mate choice, combat and courtship behaviors, species recognition, inhibition of further mating by conspecifics, and trailing (see Mason & Parker, 2010 for more details). *E. murinus* males, for example, likely locate females by following pheromone trails left by the female, using chemosensory cues to track them and distinguish between sexes (Rivas et al., 2007b). Kubie et al. (1978) found that female garter snakes (*Thamnophis radix*) treated with oestradiol benzoate became more attractive to males only after shedding their skin. Oestradiol benzoate and shedding alone were insufficient to stimulate male courtship behavior to the same extent as in treated, shedding snakes (Kubie et al., 1978). Even though oestradiol benzoate played a role in stimulating sexual attractiveness, according to the authors, shedding may have caused the release of high concentrations of male-attracting pheromones “trapped” between the old skin and the newly formed layer (Kubie et al., 1978), making them more detectable to males. It appeared male snakes assessed a female’s reproductive status by detecting hormonally regulated pheromones (Kubie et al., 1978) present in the skin.

*E. murinus’* shedding rate seems to be influenced by the snakes’ biology – including hormonal factors – rather than the growth rate of each individual (Lamonica et al., 2007). Similarly, in female timber rattlesnakes (*Crotalus horridus*), shedding events have been linked to reproductive condition, with non-pregnant females shedding around the climax of courtship behaviors (Carnes-Mason & Beaupre, 2023). If a similar reproductive-shedding relationship exists in *E. murinus*, ecdysis and mating – along with the associated courtship behaviors – may be more closely connected than previously recognized. Although the anacondas in this study were never observed copulating, the intertwining behavior exhibited may resemble the mating balls this species naturally forms during breeding (Rivas, 2007; Rivas et al., 2007b). As such, the peak in social behaviors during Phase 2 may indicate a potential link between shedding and reproduction. Nevertheless, further research is needed to determine whether ecdysis directly influences mating behavior in *E. murinus*.

Ecdysis may not only correlate with increased social behaviors but also contribute to the reduction of other behavioral categories during Phase 2. For instance, King and Turmo (1997) found that common garter snakes (*Thamnophis sirtalis*) in the shedding stage exhibited higher latencies to move beyond a delimited circle boundary after being kept under a light-blocking cover for two minutes. Among other possibilities, the researchers proposed that a movement decrease may help snakes conserve energy for the shedding process (King & Turmo, 1997). Indeed, a more recent study with timber rattlesnakes suggested that ecdysis is a highly energy-demanding process (Carnes-Mason et al., 2024), therefore, the overall decrease in locomotion, exploration, and climbing behaviors observed in this study during Phase 2 may be linked to the skin-shedding process, indicating that the anacondas may have been saving energy for other physiological demands. In the female, energy conservation was likely directed toward skin shedding, while in the male, it may have been allocated to reproduction, as previously discussed. Although not statistically significant, ecdysis may also explain the boost in body maintenance behaviors during this period (*Figure S2b*), as rubbing against objects facilitates the removal of old skin (Carnes-Mason et al., 2024). Additionally, the moderate increase in body maintenance behaviors during the final phase of the study, compared to baseline (Phase 1), suggests that enclosure modifications may have better accommodated the anaconda’s natural behavioral needs by providing more resources for the expression of rubbing behavior.

The terrarium modifications resulted in a significant increase in locomotor activity, exploration, and interaction with enhancement items during Phase 3, but not in Phase 2. This outcome was unexpected, as we anticipated a greater impact in Phase 2, while expecting activity levels to decrease in Phase 3 as the anacondas acclimated to the enclosure changes, and a decline in interest would be anticipated. The lack of research regarding *E. murinus* activity patterns in the wild, specifically the time the species allocates to each activity, makes interpreting these results challenging. Moreover, as with the previously mentioned case of inactivity, increased activity can have different implications depending on various factors, such as the species, environmental context, type of behaviors exhibited, as well as their duration and frequency. Higher activity levels may indicate improved well-being, associated with exploration and engagement with enrichment, but they may also signal stress or escape attempts. Indeed, hyperactivity has been identified as a possible abnormal behavior in captive reptiles (Warwick, 1990) and is acknowledged as a behavioral indicator of stress (Warwick et al., 2013). Warwick et al. (2013, pp. 41) described signs of hyperactivity as “abnormal high-level physical activity, surplus, or redundant activity”, none of which were observed in this study. Moreover, inactivity levels in Phase 3 did not differ significantly from those in Phase 1, and the observed behavioral variation was more indicative of behavioral diversification rather than hyperactivity. This is supported by the types of behaviors exhibited – namely, increased locomotion, exploration, and interactions with enhancement items – along with a significant yet controlled increase in these behaviors, without signs of exaggerated response or repetitive patterns. Moreover, since the terrarium had unchanged areas where the anacondas could choose to avoid the enriched sections if they preferred, and the fact that they began using the entire exhibit more evenly (*Figure 5*), suggests this is not the case and that the snakes might have been moving into these enhanced areas intentionally. Taken together, these findings highlight the importance of conducting long-term studies when assessing the impact of enclosure enhancements, allowing animals sufficient time to process the changes made, as the complete understanding of their responses may only emerge after longer adjustment periods.

In reptiles, interaction with transparent boundaries has previously been recognized as an indicator of stress in captive environments (Warwick et al., 2013). This interaction may result from living in an inadequate environment, causing the animals to actively search for alternative conditions that better meet their biological needs, or it could stem from the reptiles’ innate instincts, which, despite the transparent barrier being impenetrable, lead them to believe it can still be crossed (Warwick, 1990). According to Warwick et al. (2013), for this behavior to be considered a sign of stress, it must be persistent, occurring throughout the whole activity period. In this study, however, this behavior was brief and occasional, rather than consistent across observations, failing to meet this criterion. As such, although the anacondas exhibited a non-significant increase in interactions with the enclosure’s transparent boundaries across the three phases, the frequency of these interactions was very low, suggesting that this behavior was not a persistent sign of stress. Nonetheless, it is an aspect that should be monitored closely.

### 4.2. Enclosure Use and Environmental Parameters

Before enclosure modifications (Phase 1), both anacondas primarily used the inner lake and central terrarium regions, while showing minimal presence in the enclosure’s peripheral areas – specifically the outer lake and the terrarium’s lateral sections. These usage patterns were employed to plan the enhancements made to the terrarium, strategically focusing improvements on the less-visited areas to encourage the anacondas to explore more of their enclosure.

Phase 2 marked a shift in the anacondas’ space occupation, with the male anaconda increasing its presence in enhanced areas: the terrarium’s left side and the outer lake. In contrast, the female anaconda remained in the water throughout the entire phase. As previously mentioned, at that time, the female was undergoing ecdysis, a process where relative humidity plays a crucial role in skin shedding, as dry conditions often lead to complications during the process (Hoehfurtner et al., 2021; Carnes-Mason et al., 2024). Even though the female spent most of the observations in the inner lake, rarely using the enhanced areas, it did utilize the enhanced outer lake during the final phase of shedding, likely by moving against the trunk inside the lake, where it shed its old skin (*Figure S3*). Snakes typically use rough, uneven surfaces to aid skin shedding by moving or rubbing against these surfaces to loosen and remove the old skin (Lillywhite, 1989). The presence of shed skin on a habitat enhancement item highlighted how enhanced areas supported natural behaviors and emphasized the importance of considering an animal’s physiological state when interpreting behavioral data, since natural processes may influence terrarium preferences and space utilization in captive environments.

The final phase of this study exhibited the greatest diversity of zones used by both anacondas and in a more balanced way, particularly by the male (*Figure 5*). During this phase, there was a notable increase in the use of the terrarium’s right area, which until then had rarely been used by either anaconda. These findings suggest that terrarium modifications may have encouraged the anacondas to use a broader range of areas within their environment. In this study, the 15-day interval between Phases 2 and 3 was crucial for observing differences in the snakes’ space utilization. Skipping this adjustment period between phases could have hindered the detection of significant changes in the animals’ interaction with their environment, potentially resulting in misinterpretations of the effectiveness of enclosure modifications and the anacondas’ responses to them.

During periods of inactivity, the anacondas exhibited a significant preference for elevated surfaces in Phases 1 and 2 (*Figure 7*), likely due to the proximity of these areas to heat sources within the terrarium. Notably, the anacondas’ enclosure utilization pattern shifted dramatically in Phase 3, where they began to occupy all three heights of their enclosure – a behavior that was absent in previous phases. This more balanced utilization of the enclosure suggests that the implemented environmental enhancements may have created a more complex and stimulating enclosure, encouraging the use of previously underutilized areas. The higher occupancy of middle and lower surfaces during Phase 3 further emphasized the importance of providing animals with adequate time to adjust to changes in their environment.

While significant temperature gradient shifts occurred between Phases 1 and 2 (*Figure 8*), no significant differences were detected between Phases 2 and 3, suggesting that the behavioral and spatial differences observed in the final phase likely resulted from terrarium modifications rather than temperature fluctuations. However, the relationship between behavioral patterns, space utilization and habitat enhancements requires cautious interpretation, as significant temperature variations were recorded across all phases on both the left and right sides of the terrarium (*Figure 8*). Additionally, we identified a strong negative relationship between temperature and relative humidity – both critical factors for reptile behavior (Daltry et al., 1998; Sunday et al., 2014) – indicating that the observed changes in behavioral patterns and enclosure use may have been influenced by these environmental variables rather than terrarium modifications alone. Nonetheless, the temperature shift was more pronounced between Phases 1 and 2 (≈0.9 ºC and ≈2 ºC on the left and right sides, respectively) than between Phases 2 and 3 (≈0.3 ºC and ≈0.6 ºC on the left right sides, respectively), which further reinforces the likelihood that behavioral and spatial changes were related to enclosure modifications than to thermal conditions.

### 4.3. Limitations

This study had a few limitations that, even though we consider do not detract from the validity of our findings, should be acknowledged. The scarcity of published research on *E. murinus*, specifically regarding its behavioral repertoire and time allocation patterns, made it challenging to fully interpret our results, as few studies were available for comparison. Additionally, conducting research with animals in captivity presented the inherent limitation of a restricted sample size. Other variables, such as sex differences and the ongoing ecdysis process in one of the individuals, may have also contributed to variations in the results, particularly during Phase 2. To build on these findings, future research would benefit from extended observation periods and a larger number of observations. As such, even though this study provides meaningful insights into the species’ behavior and management under human care, these methodological constraints should be considered when interpreting our results.

## 5. Conclusion

The findings from this study indicate that enclosure enhancements positively influenced the anacondas’ behavior by increasing behavioral diversity, promoting more balanced space utilization, encouraging exploration of the enclosure’s vertical dimension, and leading to the occupation of previously underutilized areas. Curiously, the anacondas demonstrated increased interaction with terrarium enhancement items during the study’s third and final phase. The interval between Phases 2 and 3 provided valuable insights into the extended effects of enclosure modifications, demonstrating the importance of allowing sufficient time for the anacondas to adjust to their new environment and ensuring that both short and long-term responses to the enhancements were captured.

## Author Contributions

Conceptualization, M.I.P. and A.M.; Methodology, A.M.; Validation, M.I.P; J.C.; G.M.; F.R. and A.M.; Formal Analysis, G.M. and A.M.; Investigation, M.I.P; Resources, F.R.; Writing – Original Draft Preparation, M.I.P and J.C.; Writing – Review & Editing, M.I.P; J.C.; G.M.; F.R. and A.M.; Visualization, M.I.P.; J.C.; G.M. and A.M.; Supervision, F.R. and A.M.; Project Administration, F.R.; A.M.; All authors have read and approved the submitted version and agree to be accountable for the contributions and for ensuring that questions related to the accuracy or integrity of any part of the work are appropriately investigated, resolved, and documented in the literature.

## Institutional Review Board Statement

The study was conducted according to the guidelines of the Declaration of Helsinki, and approved by the Animal Welfare Committee of ICBAS, School of Medicine and Biomedical Sciences of the University of Porto (P480/2023/ORBEA, 4^th^ November 2024).

## Informed Consent Statement

Not applicable.

## Data Availability Statement

The raw data supporting the conclusions of this article will be made available by the authors upon reasonable request.

## Acknowledgements

We thank Zoo Santo Inácio for its continual support and exceptional accessibility throughout this research. A special thank you to the entire zoo team for their contributions, both direct and indirect, from those who implemented the enclosure enhancements to everyone who participated in all the valuable discussions.

