## Supplementary Materials for "Give It Time: Green Anacondas Show Prolonged Behavioral Responses to Enclosure Enhancements"

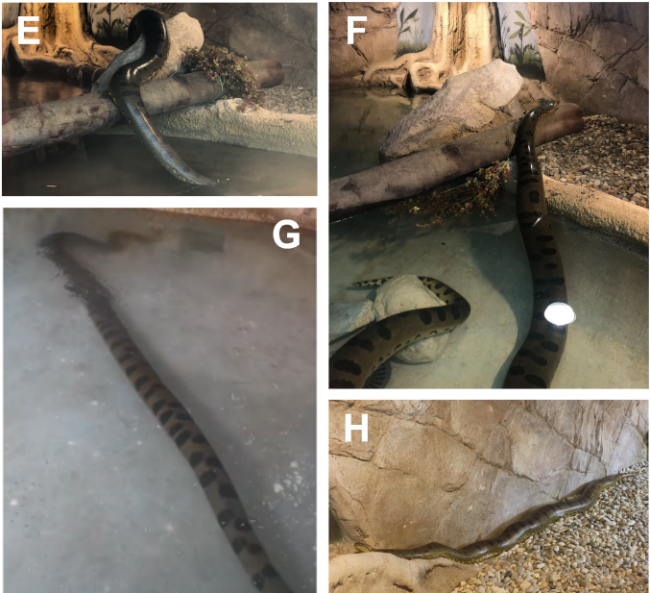

**Figure S1 (cont).** Ethogram of the green anacondas (*Eunectes murinus*) observed in this study. **LOCOMOTION (E-H):** **Entering the lake (E) –** Movement from land to water (headfirst or by moving backward), from initial contact until half of the body is submerged. Beyond this point, the subject is in the water rather than entering it. **Exiting the lake (F) –** Movement from water to land, until half the body has emerged. **Swimming (G)** **–** Movement through the water using lateral undulations while in the water column, either submerged or with its head at the surface. **Slithering (H) –** Movement that occurs on land or in water through lateral undulations, with the ventral side contacting the ground. On land, the head may be elevated a few centimeters, while in water, it can be submerged, with the head lifted from the lake’s bottom or at the surface. It may involve partial or full-body movement.

**Figure S1.** Ethogram of the green anacondas (*Eunectes murinus*) observed in this study. **INACTIVE (A-D): Resting submerged in water (A-C) –** Remaining motionless in water, submerged and usually resting at the bottom of the lake, or with the head near the surface, either completely above the water, with only eyes and nose exposed, or with the anterior half of the body exposed. The body may be coiled or stretched out. **Resting on land (D) –** Remaining motionless on land, fully out of the water. The body may be coiled with the head resting on top or stretched out. Occurs at various heights within the enclosure.

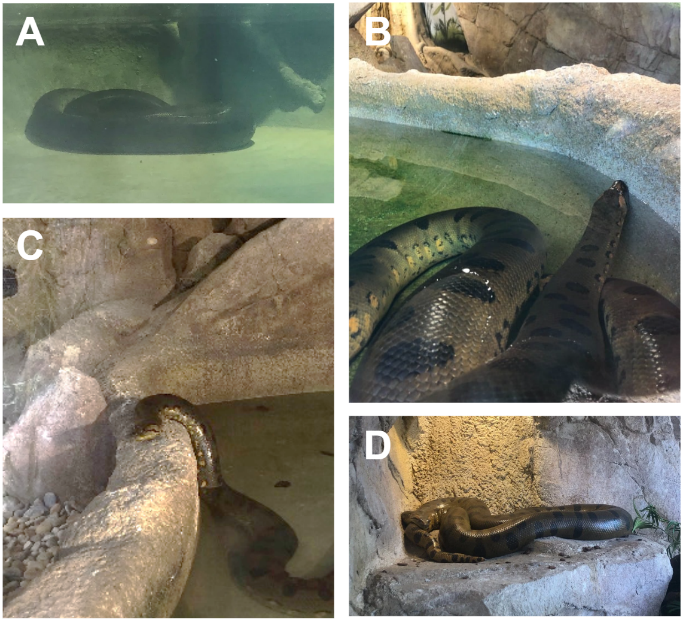

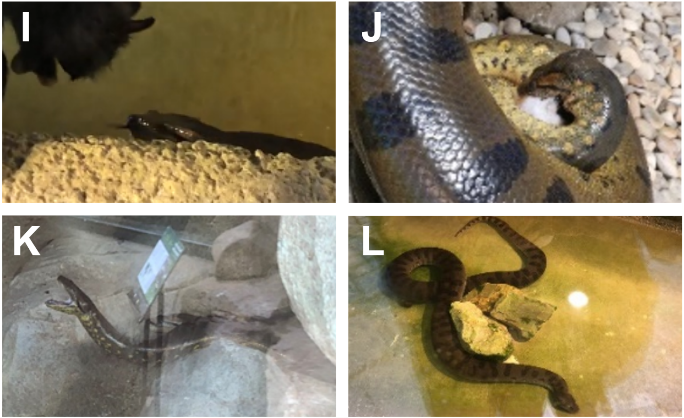

**Figure S1 (cont).** Ethogram of the green anacondas (*Eunectes murinus*) observed in this study. **FEEDING (I-J): Food pre-capture (I) –** Intent staring at prey, moving toward it, often tongue flicking (see below). **Food capture (J) -** Attacking, dragging the prey to water (if not on land), and constricting it. **BODY MAINTENANCE (K-L): Yawning (K) –** Brief mouth opening (3- 5s) while inactive, not directed at individuals or objects. **Rubbing (L) –** Moving the body’s side against an object, either on land or in water.

**Figure S1 (cont).** Ethogram of the green anacondas (*Eunectes murinus*) observed in this study. **INTERACTION WITH TRANSPARENT BOUNDARIES (O-P): Crawl up the high window (O) –** Lifting at least one-third of the anterior body against the glass, appearing to attempt to climb it. It may occur on land or in water. **Swimming against the high window (P) –** Moving the head against the glass while in water.

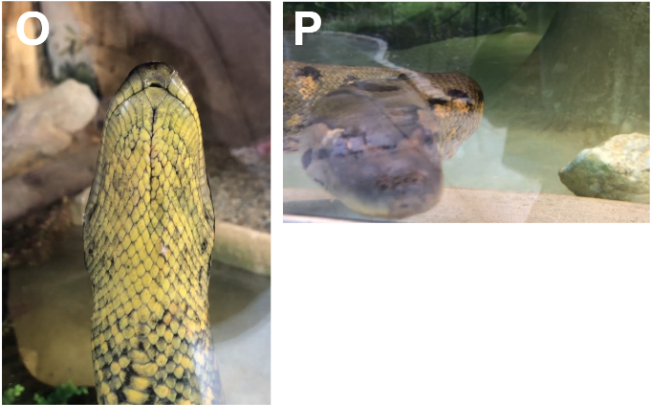

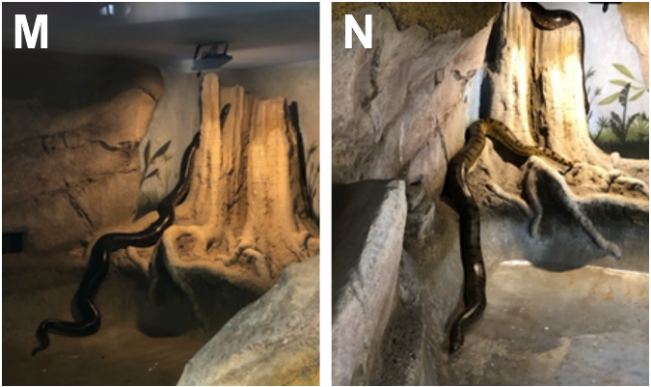

**Figure S1 (cont).** Ethogram of the green anacondas (*Eunectes murinus*) observed in this study. **CLIMBING (M-N): Climbing up (M) –** Ascending to higher surfaces by undulating using undulations with more than half the body engaged. **Climbing down (N) –** Descending to lower surfaces using undulations with more than half the body engaged.

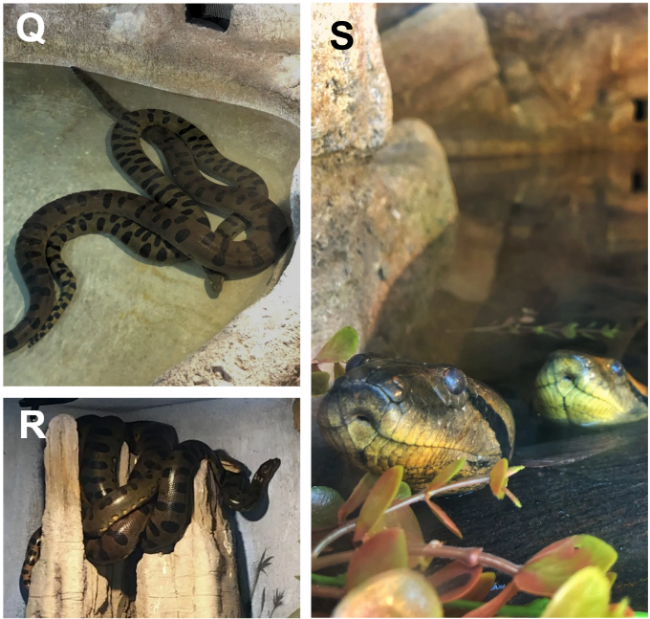

**Figure S1 (cont).** Ethogram of the green anacondas (*Eunectes murinus*) observed in this study. **SOCIAL INTERACTION^1^ (Q-S): Coiling around conspecific (Q-R)** – Intertwining the body around a conspecific. It may occur at various heights, on land or in water. **Proximity to a conspecific (S) –** Remaining near another anaconda (< 50 cm) without physical contact, on land or in water.

^1^ Can occur simultaneously with other behavioral categories. For instance, an anaconda may remain inactive or move while coiled around a conspecific.

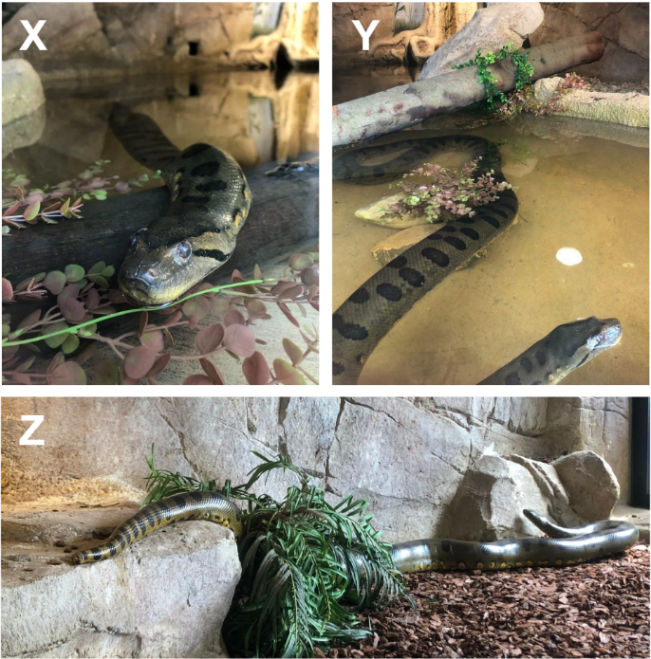

**Figure S1(cont.).** Ethogram of the green anacondas (*Eunectes murinus*) observed in this study. **ENGAGING WITH HABITAT ENHANCEMENT ITEMS3 (X-Z):** **Resting on habitat enhancement items (X) –** Remaining inactive while positioned on top of habitat-enhancing items, either on land or in water. **Rubbing against habitat enhancement items (Y) –** Moving the side of the body against habitat-enhancing items, on land or in water. **Slithering on habitat enhancement items (Z) -** Moving over habitat-enhancing objects, whether on land or in water. **NOT VISIBLE:** The anaconda is out of sight.

^3^ Can occur simultaneously with other behavioral categories. For example, an anaconda may rest on environmental enhancement items while near a conspecific or may actively explore these items.

**Figure S1 (cont).** Ethogram of the green anacondas (*Eunectes murinus*) observed in this study. **EXPLORATION (T-W): Tongue flicking^2^ (T) –** Extending, flicking, and retracting tongue. **Head or body elevation (U-V) –** Raising at least one-third of the anterior body vertically or horizontally, while stationary. The head may remain aligned with the body or parallel to the ground. It may include tongue flicking or body undulations, and be directed at an object or individual, occurring both in and out of the water. **Head or body lowering (W) –** Lowering at least one-third of the anterior body while stationary. It may include tongue flicking or body undulations and be directed at an object or individual.

^2^ Can occur simultaneously with other behavioral categories. For example, an anaconda may rub against an object or slither through the terrarium while flicking its tongue.

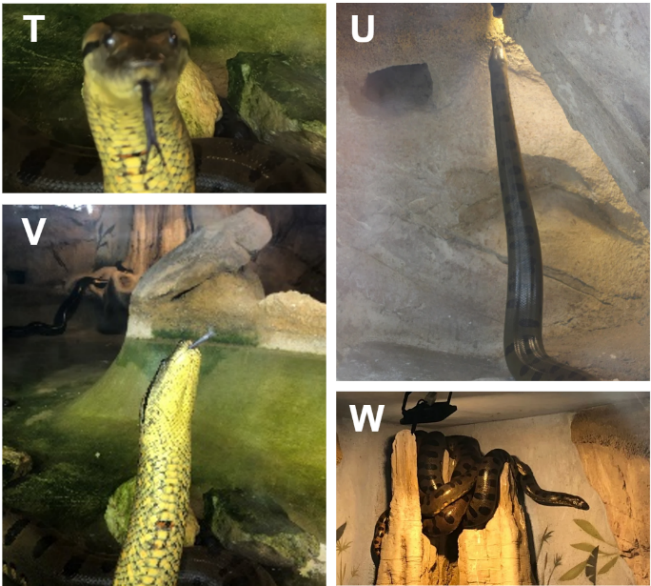

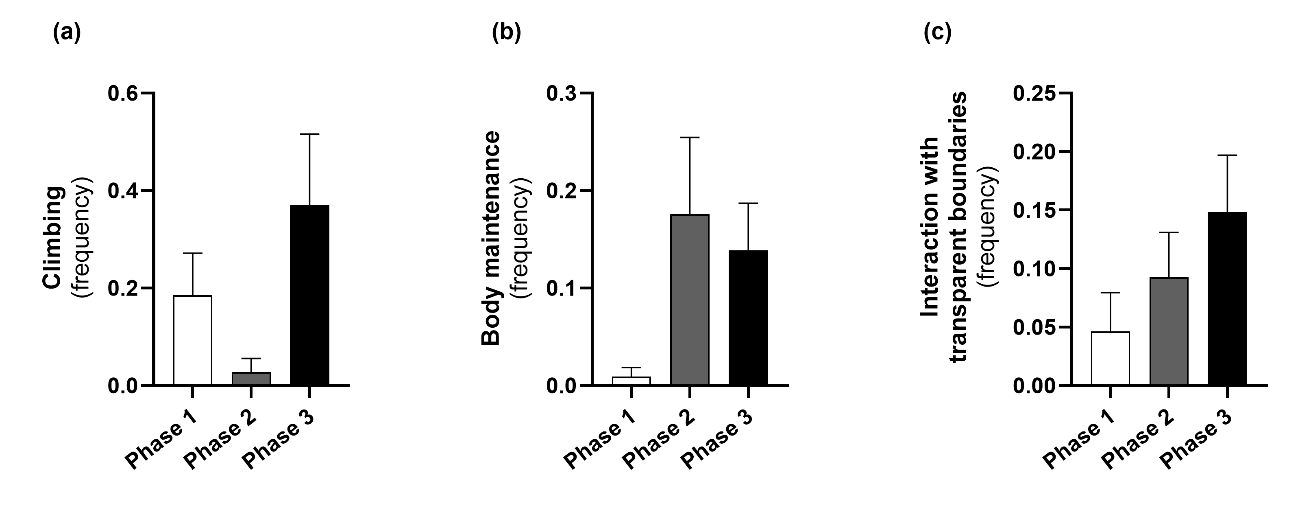

**Figure S2.** E. murinus behavioral changes to habitat enhancement across the three study phases. Panels display the frequency of **(a)** climbing, **(b)** body maintenance, and **(c)** interactions with transparent boundaries. **Phase 1 –** Before enclosure modifications; **Phase 2 –** Immediately after enclosure modifications; **Phase 3 –** Approximately three weeks after enclosure modifications. Data is represented as mean ± SEM.

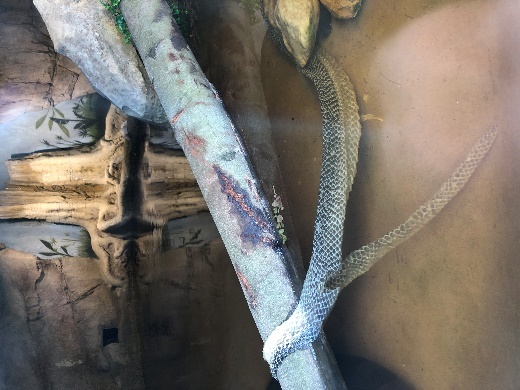

**Figure S3.** The female anaconda shed skin during Phase 2. The old skin was left on an enhanced area (outer lake), likely by moving against the tree trunk.
